# Elucidating cryptic sympatric speciation in terrestrial orchids

**DOI:** 10.1101/828491

**Authors:** Emerson R. Pansarin, Alessandro W.C. Ferreira

**Affiliations:** University of São Paulo, FFCLRP-USP, Department of Biology, Av. Bandeirantes 3900, 14040-901, Ribeirão Preto, SP, Brazil; Federal University of Maranhão, Campus Dom Delgado, Department of Biology, Av. dos Portugueses, 1966, 65080-805, São Luís, MA, Brazil

**Keywords:** Epidendroideae, mesophytic semideciduos forest, Orchidaceae, reproductive biology, reproductive isolation

## Abstract

**PREMISE:** Cryptic sympatric speciation occurs when closely related species arises with no geographic or spatial isolation. Since cryptic species can not usually be detected when investigations are based exclusively on classical plant taxonomy, molecular markers and integrative taxonomy are important tools to elucidate the identity of cryptic taxa.

**METHODS:** In this paper we aimed to provide a detailed investigation based on molecular data and experimental taxonomy conducted on populations of a plant model occurring inside mesophytic forests and a supposed cryptic sympatric species that is found in marshy areas. Furthermore, a phylogeny was reconstructed by using nrDNA and cpDNA sequences in order to ascertain the position of both sympatric species within *Liparis*. The divergence among sequences of the putative cryptic sympatric species and related taxa was also verified based on DNA barcoding.

**KEY RESULTS:** Our results reveal cryptic sympatric speciation in the orchid genus *Liparis*. Differences in microhabitat, flowering phenology, morphology of leaves and flowers, and reproductive strategies support the taxon occurring in marshy areas is a cryptic species. The molecular analyses reveal the cryptic species is closely related to *L. nervosa*.

**CONCLUSIONS:** Paludal *Liparis* is a sympatric cryptic species distinguishable from *L. nervosa* by differences in habitat, phenology and breeding system. Our conclusions are supported by the molecular investigations. The cryptic *Liparis* emerges as sister to *L. nervosa*, within a clade with terrestrial habitat and plicate leaves. The discovery of sympatric cryptic speciation in orchids provides increase on the knowledge of diversification and reproductive isolation in flowering plants.

---

Cryptic species is when two or more taxa that have been considered as a unique nominal taxon are practically indistinguishable when based solely on morphological characteristics (Bickford et al., 2007). Many studies have shown that the occurrence of cryptic species is relatively common, but many of them cannot be detected when investigations are based exclusively on classical plant taxonomy (i.e. morphology). The use of molecular markers and integrative taxonomy, which includes inter-specific crosses, breeding data, investigations on chemistry, cytology and pollinators plus floral biology have been used to elucidate the identity of cryptic plant species (Stuessy, 2007). More recently, the use of DNA sequencing has been helpful in species identification and in recognizing cryptic species, since sequence divergence is usually lesser between individuals of a species than among closely related taxa (Herbert et al., 2004). Cryptic species occur in many groups of animals (Herbert et al., 2004; Priti et al., 2016), but also have been recorded in plants (Hollingsworth et al., 2009), including orchids (Pansarin and Amaral, 2006; Pace et al., 2017; Gale et al., 2018).

Sometimes species arises with no geographic or spatial isolation, occurring as sympatric, but reproductively isolated (Bolnick and Fitzpatrick, 2007). This process has been referred as sympatric speciation (Mayr, 2000). Cryptic sympatric speciation has been reported for a number of organisms, but rarely has been recorded in plants (Les et al., 2015). At least is known, cryptic sympatric speciation never be recorded in Orchidaceae. In orchids, pollinator-mediated sympatric speciation has been commonly referred to species pollinated by sexual deception (Xu et al., 2012). In fact, orchids are considered good plant models to investigate sympatric diversification, since the pollinators pressures result in shifts on flower morphology and floral fragrance that are easily identified by usual methods in plant taxonomy (Xu et al., 2012). However, in the case of cryptic speciation, often these morphological markers are obscure and pre-pollination barriers commonly are weakly or absent. Furthermore, is hard to verify reproductive isolation in plants, since lineages can differ by polyploidization, and an incomplete process of reproductive isolation can result in post-polyploid introgression. Additionally, only a few phylogenetic inferences include samples from more than one population in the analyses. As a consequence, a number of cryptic plant species could stay undetected as a consequence of an inadequate methodology or poor sampling (Les et al., 2015). So, if cryptic sympatric species seems to be common in plants, it also can be detected in orchids? In this paper we have selected populations of terrestrial species of *Liparis* as model to try to answer this question.

*Liparis* Rich. (Malaxideae) is a cosmopolitan genus that comprises more than 350 species occurring as terrestrial or epiphytes throughout temperate and tropical regions of the world (Pridgeon et al., 2005). The genus is characterized by soft and sheathing leaves, frequently with plicate leaf blades, ovoid and often subterranean bulb, and raceme with resupinate flowers. It has been considered polyphyletic, since at least two genera (i.e. *Malaxis* and *Oberonia*) emerge within *Liparis* (Cameron, 2005; Tsutsumi et al., 2007). In *Liparis* there are three main groups: one clade whose species are predominantly epiphytes, and two terrestrial clades, one of them including species with conduplicate leaves and another that comprises species with flat leaf blades (Cameron, 2005; Tsutsumi et al., 2007). In Brazil occur three species of *Liparis*: *L. cogniauxiana* F. Barros & L. Guimarães (=*L. bifolia* Cogn.), *L. nervosa* (Thunb.)Lindl. and *L. vexillifera* (La Llave & Lex.) Cogn.(Barros et al., 2019). During the developing floristic surveys of the orchid family in the state of São Paulo, Brazil (Pansarin and Pansarin, 2008, 2010; Ferreira et al., 2010), populations of an unidentified *Liparis* occurring in marshy open areas have been found. This species possesses vegetative and floral characteristics closely related to those of *L. nervosa*, a species that occur in the forest understory, and the distinction of both taxa based exclusively on morphological characteristics from herbaria material is obscure. As consequence, this taxon has been considered as *L. nervosa* in floristic inventories (Pansarin and Pansarin, 2008, 2010; Ferreira et al., 2010). Nowadays, evidences that this marshy Liparis could be a sympatric cryptic species has been materializing, since plants collected in the field and cultivated under the same conditions show differences with regards the phenology, details in vegetative and floral morphology and fruit set. Based on the above concerns, a detailed investigation based on living plants and on experimental taxonomy was carried out on populations of *L. nervosa* and this potential sympatric cryptic taxon. Furthermore, a phylogenetic inference was made by using nrDNA and cpDNA sequences in order to determine the position of members of both sympatric populations within *Liparis*. In addition, the divergence among sequences of this putative sympatric cryptic species and related taxa was based on DNA barcoding analysis and their relationships discussed.

## MATERIALS AND METHODS

### Study site and plant material

Our study was based on plants growing in the natural reserve of Serra do Japi, municipality of Jundiaí (approx. 23°11’S, 46°52’W; 910 m a.s.l.), state of São Paulo, southeastern Brazil. This region is characterized primarily by mesophytic semi-deciduous forests (Leitão-Filho, 1992).

The experiments were based on specimens of *Liparis* collected in open marshy areas and inside the forest. Ten adult plants of *Liparis nervosa* and 15 plants of *Liparis* sp. growing at least 30 m away from each other were collected from the natural population and planted with coconut fiber in plastic pots. Plants were maintained under the same irrigation and light (50% black shade cloth) conditions. The plants were collected in February 2013 and grown at the LBMBP Orchid House, University of São Paulo. The LBMBP Orchidarium is entirely covered in 50% black shade cloth and isolated from the outside to avoid the contact of the plants with any possible flower visitor. The LBMBP Orchid House is located in the municipality of Ribeirão Preto (approx. 21°10’S, 47°48’W; 546 m a.s.l.) ca. 250 km NW away from the study area. Ribeirão Preto and Jundiaí are situated in an ecotonal area between Cerrado and Atlantic Forest (Kronka et al., 1993) where the climate is classified as “Cwa” (i.e. mesothermic with a dry winter season; Köppen, 1948).

One specimen of both studied species was vouchered and deposited at the Brazilian herbaria, as follow: *Liparis nervosa*: Brazil, São Paulo, Jundiaí, Serra do Japi: 14.XII.1997, *E.R. Pansarin 100* (UEC); *Liparis* sp.: Brazil, São Paulo, Jundiaí, Serra do Japi: 16.II.2017, *E.R. Pansarin 1544* (LBMBP). The plant species were identified based on floristic inventories performed in the state of São Paulo (Pansarin and Pansarin, 2008, 2010; Ferreira et al., 2010).

Fieldwork was performed according the Brazilian legislation. Permission by Base Ecológica da Serra do Japi, and SISBIO (process number 35178-1).

### Flowering phenology, plant features and spontaneous self-pollination experiments

Flowering phenology and flower lifespan of both *Liparis nervosa* and *Liparis* sp. were recorded by visiting the study site in Serra do Japi monthly, from February 2012 to March 2016.

The morphological differences between *Liparis nervosa* and the putative cryptic species occurring in marshy areas (*Liparis* sp.) were recorded based on plants collected in the field and cultivated at the LBMBP Orchid House. Vegetative and reproductive structures of *Liparis nervosa* (10 plants) and *Liparis* sp. (15 plants) were studied. Floral details of *Liparis nervosa* (30 flowers; six plants; six inflorescences; five flowers per inflorescence) and *Liparis* sp. (30 flowers; six plants; six inflorescences; five flowers per inflorescence) were analyzed under a stereomicroscope Leica S8 APO. Measurements of floral structures were made with a Vernier caliper. Floral morphology was investigated concerning the shape and size of floral structures such as sepals, petals and labellum, column, anther, and pollinarium (Faegri and van der Pijl, 1979).

The investigation of spontaneous self-pollination was based on 10 plants of *Liparis nervosa* and 15 plants of *Liparis* sp. during two flowering seasons (i.e. from November 2014 to February 2015, and from November 2015 to February 2016). In the 2014-2015 flowering season, the plants, including flowers, were submitted to a unique treatment inside the Orchid House: irrigation once a day, simulating rain. In the 2015-2016 flowering season, these same plants were submitted to two treatments: seven plants of *L. nervosa* and 12 plants of *Liparis* sp. were maintained inside the Orchid House and then irrigated (including the flowers) once a day, simulating rain. The remaining six plants (three plants of *Liparis nervosa* and three plants of *Liparis* sp.; two inflorescences per plant) were placed in three glass boxes (60 × 50 × 40 cm each), which were maintained inside the LBMBP Orchidarium, under the same conditions of the preceding tests, but without irrigating the inflorescences. Thus, each box contained one plant (two inflorescences) of *Liparis* sp. and one plant (two inflorescences) of *L. nervosa*. The bottom of the boxes remained opened during the experiment for airflow and watering of the roots and substrate. The occurrence of spontaneous self-pollinations was determined by counting the number of fruits formed in relation to the total of flowers produced per inflorescence. The fruit set was recorded when fruits were dehiscent.

Additionally, to determine the influence of environmental conditions (e.g. rainfall and air humidity) on spontaneous self-pollination, the temperature and the relative humidity were monitored hourly during the flowering period of both *Liparis nervosa* and *Liparis* sp., from 13 November 2015 to 15 February 2016. The temperature and the relative humidity were recorded with an Extech *RHT10* - Humidity and Temperature USB *Data Logger*. The temperature/humidity data logger was installed inside one of the three glass boxes, in the LBMBP Orchid House, FFCLRP-USP.

### Taxon sampling for phylogenetic analysis

To establish the phylogenetic position of the putative cryptic *Liparis*, a matrix containing sequences of Malaxidae genera (i.e. *Liparis, Acanthephippium* Blume ex Endl., *Collabium* Blume, *Crepidium* Tausch, *Dienia* Lindl., *Eria* Lindl., Malaxis Sol. ex Sw. and *Oberonia* Lindl.) was analyzed. The matrix was rooted with *Eria*. The list of outgroup and ingroup taxa, their GenBank accession numbers and vouchers are given in Appendix S1.

Additionally, the divergence among the putative cryptic taxon and their sister species was estimated based on DNA barcoding analyses. Two loci were amplified for each of the vouchered samples: ITS (nrDNA) and *matK-trnK* (cpDNA), which are commonly used in routine DNA barcoding (Li et al., 2011). The specimens used in DNA barcoding analysis, the vouchers, and the GenBank numbers are presented in Appendix S1.

### DNA extraction, amplification and sequencing

The DNA extraction was made from fresh leaves following an adapted CTAB method (Doyle and Doyle, 1987). The amplifications were performed using 50 μL PCR reaction. A 5 M Betaine was included to PCR reaction for DNA relaxation. Primers for the regions ITS1, 5.8S and ITS2 (Sun et al., 1994) and *matK-trnK* (Johnson and Soltis, 1995) were used for the amplifications and sequencing. *Taq* DNA polymerase was included to the PCR mix at 80 °C after a denaturation period of 10 min at 99 °C. 35 cycles were run following the programation: denaturation, 1 min, 94 °C; annealing, 45 sec, 64 °C (ITS), 54 °C (*matK-trnK*); extension, 1 min, 72 °C; final extension, 5 min, 72 °C. The PCR products were purified with GFX columns (GE Health Care). The preparation of the 10 μL sequencing reactions was made by using Big Dye 3.1 (ABI). Sequences were obtained with a 3100 Applied Biosystems sequencer. For editing the sequences assembly of complementary and overlapping sequences, the program Sequence Navigator and Autoassembler (Applied Biosystems) were used. All regions were aligned with BioEdit version 5.0.9.

### Phylogenetic analyses

The analyses of Maximum parsimony (MP) were performed with PAUP* version 4.0b5 (Swofford, 2001). Heuristic searches were run with 69 taxa for ITS, 83 for *matK-trnK*, and 69 for the combined data matrix. The heuristic searches for individual and combined data matrices were run with 1000 replicates of random taxon entry additions, tree bisection-reconnection (TBR) branch-swapping algorithm and MULTREES option, holding 10 trees per replicate and saving the shortest trees. Support for clades was assessed with 5,000 bootstrap replications (Felsenstein, 1985).

The Bayesian analysis (BI) was run with MrBayes version 3.1 (Ronquist and Huelsenbeck, 2003), while Maximum likelihood (ML) analysis of concatenated loci was run with both PAUP* version 4.0b5 (Swofford, 2001). For both BI and ML, a combined data matrix of ITS and *matK-trnK* containing 69 taxa (2919 characters) was analyzed. The data matrix based on the combination of the data was partitioned in two categories (ITS and *matK-trnK*), while the model of evolution of sequences for the partitions was selected with jModeltest (Posada, 2008), and under BIC statistics. The program selected the evolution model SYM+I+G for ITS (nrDNA) partition and GTR+G for *matK-trnK* (cpDNA) partition. In the ML analysis initial tree(s) for the heuristic search were obtained automatically by applying Heuristic Search algorithm to a matrix of pairwise distances estimated using the Maximum Composite Likelihood (MCL) approach, and then selecting the topology with superior log likelihood value. Standard non-parametric bootstrapping was employed with 1000 replicates. Bootstrap support (BS) values above 50% were mapped on branches of the tree. For BI analysis, four Markov chains were performed for three million generations, with parameters tested every 100 generations. The consensus tree was calculated after removal of the first 3,000 trees, here referred as “burn-in”. Posterior probability (PP) values above 0.5 were mapped on branches of the consensus tree. Maximum likelihood tree was draw with MEGA7 (Kumar et al., 2016), while the BI tree was accessed with Mesquite (Maddison and Maddison, 2010).

### DNA barcoding analysis

The molecular divergence among the putative cryptic *Liparis* and their sister taxon was estimated based on the barcoding regions: ITS (nrDNA), and *matK-trnK* (cpDNA). To estimate the evolutionary divergence between sequences, the number of base substitutions per site was calculated by a bootstrap procedure (1,000 replicates). Analyses were conducted using the Kimura 2-parameter model (Kimura, 1980). Codon positions included were 1st+2nd+3rd+Noncoding. All positions containing gaps and missing data were eliminated. There were a total of 3240 positions in the final dataset. The analyses were conducted in MEGA7 (Kumar et al., 2016). In addition, a tree of K2P distances was inferred using the Neighbor-Joining method (Saitou and Nei, 1987) to present a graphic illustration of the divergence pattern among the sequences.

## RESULTS

### Flowering phenology, plant features and spontaneous self-pollination experiments

Based on investigations in the Serra do Japi, *Liparis nervosa* was collected exclusively inside the forest, in well-drained soils, whereas the putative cryptic *Liparis* species was gathered solely from marshy open areas. *Liparis nervosa* flowers from November to January, while the flowering period of the putative cryptic taxon (*Liparis* sp.) occurred from January to February. This same phenological pattern was recorded in plants cultivated under the same conditions (water, temperature and luminosity) at the LBMBP Orchid House since 2013 (Fig. 1). Main differences between *Liparis* sp. and their related taxon (*L. nervosa*) are summarized in Table 1.

**FIGURE 1.**
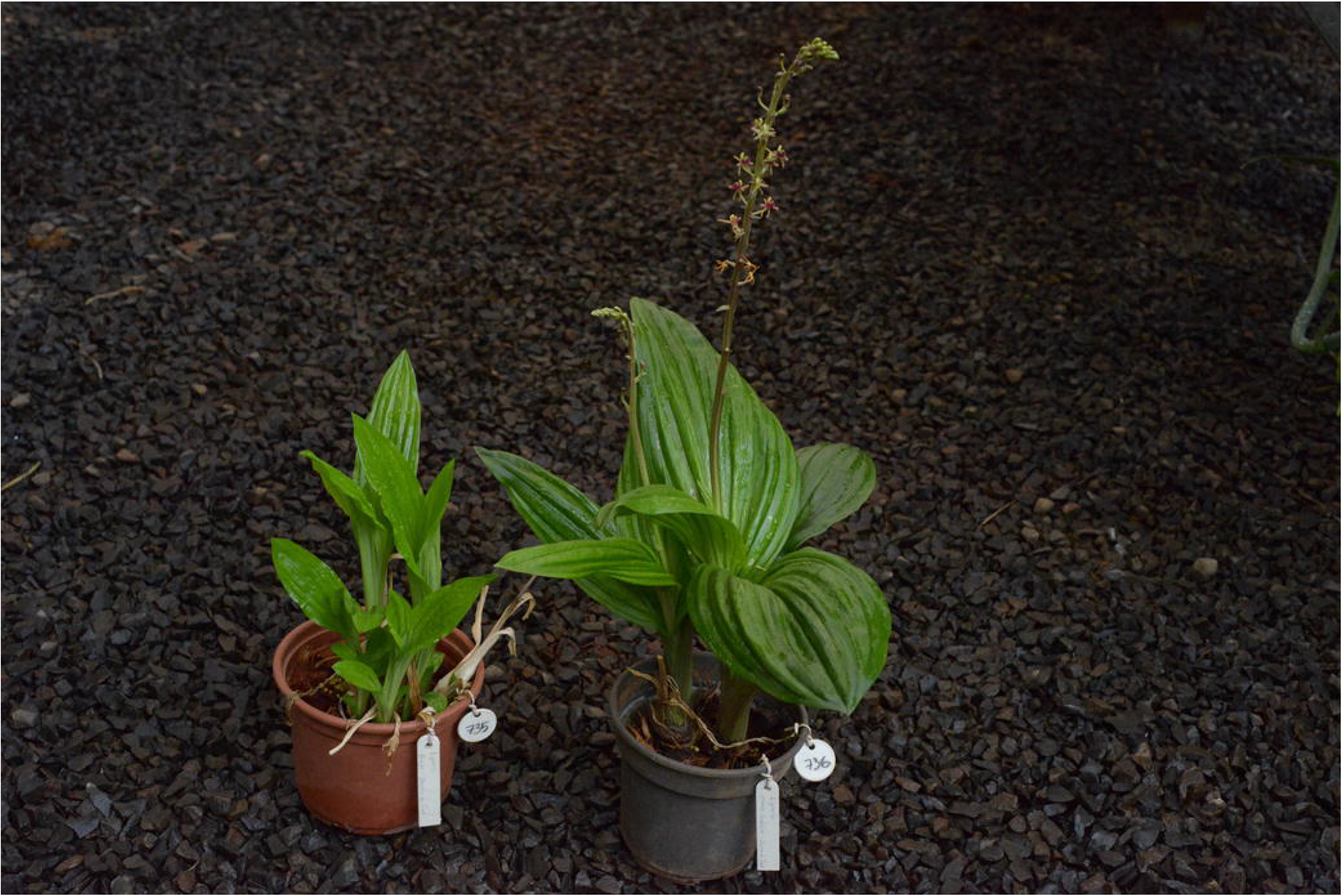
*Liparis* sp. (left) and *Liparis nervosa* (Thunb.) Lindl. (right) in cultivation at the LBMBP Orchidarium. Note *L. nervosa* with open flowers, while plants of *Liparis* sp. are in vegetative stage.

**TABLE 1.** Diagnostic morphological characters of the cryptic species of *Liparis* and *L. nervosa* from Serra do Japi, southeastern Brazil.

| Feature | *Liparis* sp. | *Liparis nervosa* |
| --- | --- | --- |
| Vegetation formation | Marshy open vegetation | Mesophytic semidecidual forest |
| Habit | Paludose | Terrestrial |
| Length of adult pseudobulbs<br>(flowering plants) | Less than 5 cm | More than 5 cm |
| Colour of adult pseudobulbs | Pale green | Purplish green |
| Width of apical leaf blades | 3.5-5.5 cm | 6-11 cm |
| Colour of the adaxial surface of<br>leaves | Pale green | Dark green |
| Colour of the abaxial surface of<br>leaves | Pale green | Purplish green |
| Colour of the inflorescence axis | Pale green | Purplish green |
| Leaf blade shape | Narrowly elliptic to<br>lanceolate | Largely elliptic to ovate |
| Colour of sepals and petals | Greenish | Purplish |
| Width of the labellum | Less than 5.8 mm | More than 6 mm |
| Labellum colour | Purplish green | Dark purple |

The treatment of spontaneous self-pollination (exposed to rainfall but pollinators excluded) produced a fruit set of 36.7% for *Liparis* sp., while no fruits were formed in flowers of *L. nervosa* (Fig. 2). In the treatment where both pollinator and rainfall were excluded, with a variation in air temperature from 16.9 to 39.8 °C, and a variation in relative air humidity from 35 to 99.9% (Appendix S2), plants of *Liparis* sp. averaged 21.9% fruit set, while no fruits were formed for *L. nervosa* (Fig. 3). Thus, our results revealed that rainfall is unnecessary for the occurrence of spontaneous self-pollination in *Liparis* sp. (Fig. 4). Furthermore, no fruits were formed in plants of *Liparis nervosa*, suggesting a biotic agent is needed for pollination (Figs. 2–4). Autogamy was recorded when the viscidium remained attached to the rostellum while the pollinia bended down and contacted the stigmatic surface. During this process, the anther cap remained attached to the apex of the column.

**FIGURE 2.**
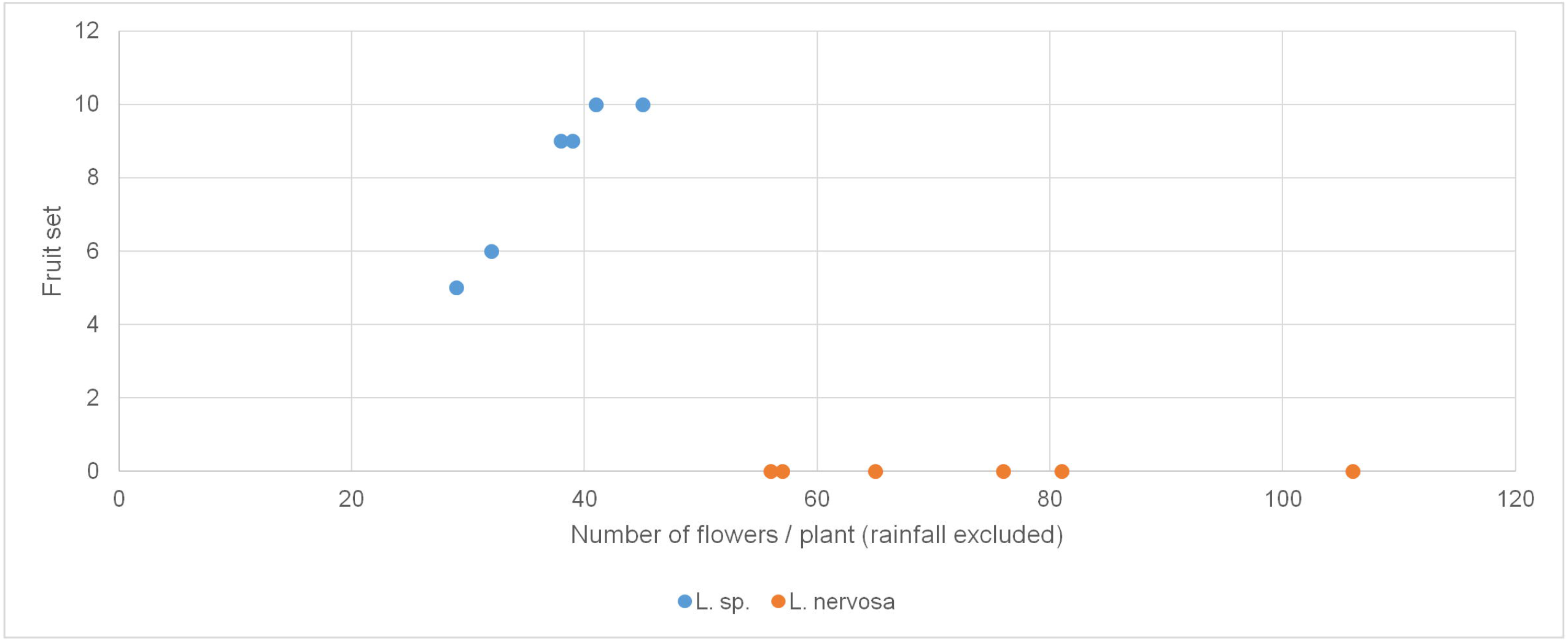
Fruit set in relation to the number of flowers produced per plant in individuals of *Liparis nervosa* and *Liparis* sp. collected in populations of the Serra do Japi and maintained under glass boxes (rainfall and pollinators excluded).

**FIGURE 3.**
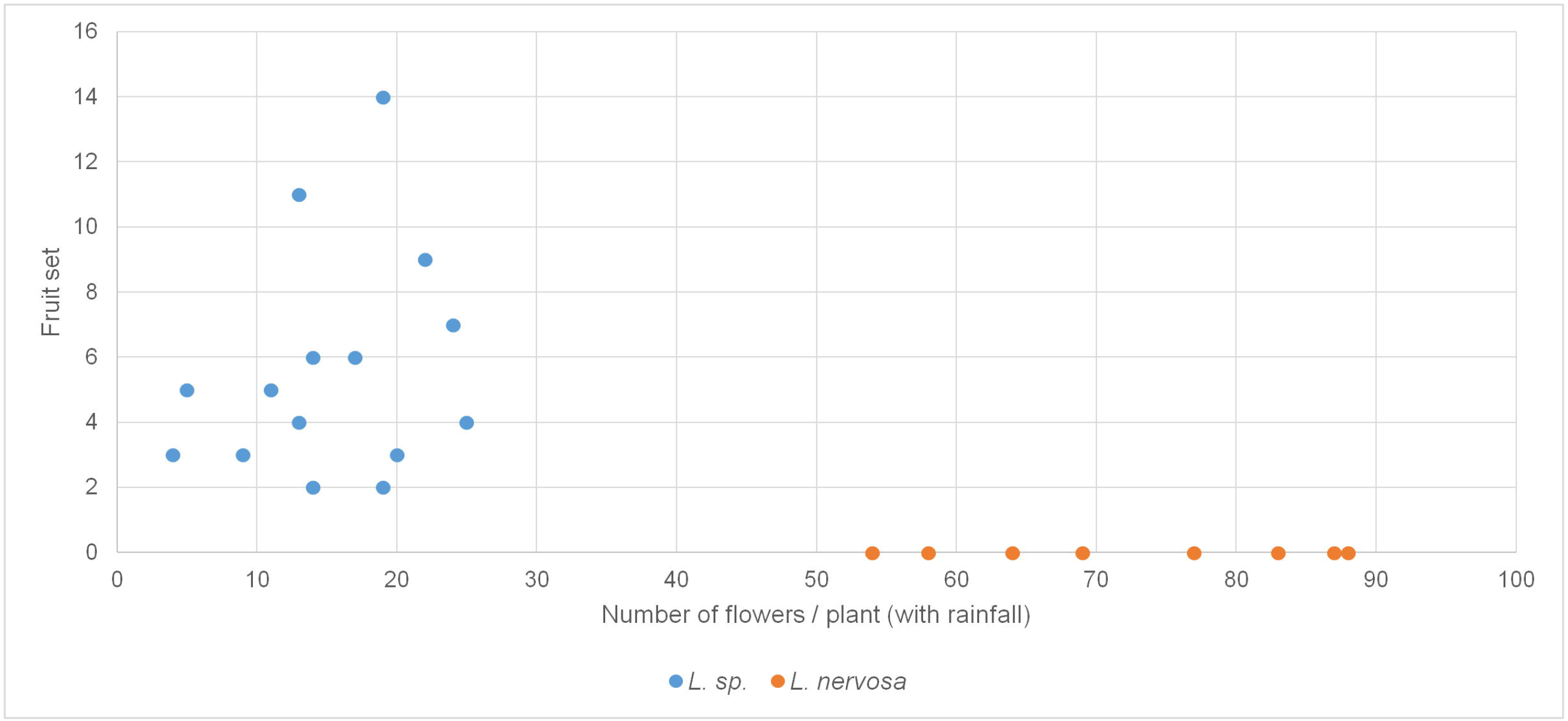
Fruit set in relation to the number of flowers produced per plant in individuals of *Liparis nervosa* and *Liparis* sp. collected in populations of the Serra do Japi (pollinators excluded, but with rainfall).

**FIGURE 4.**
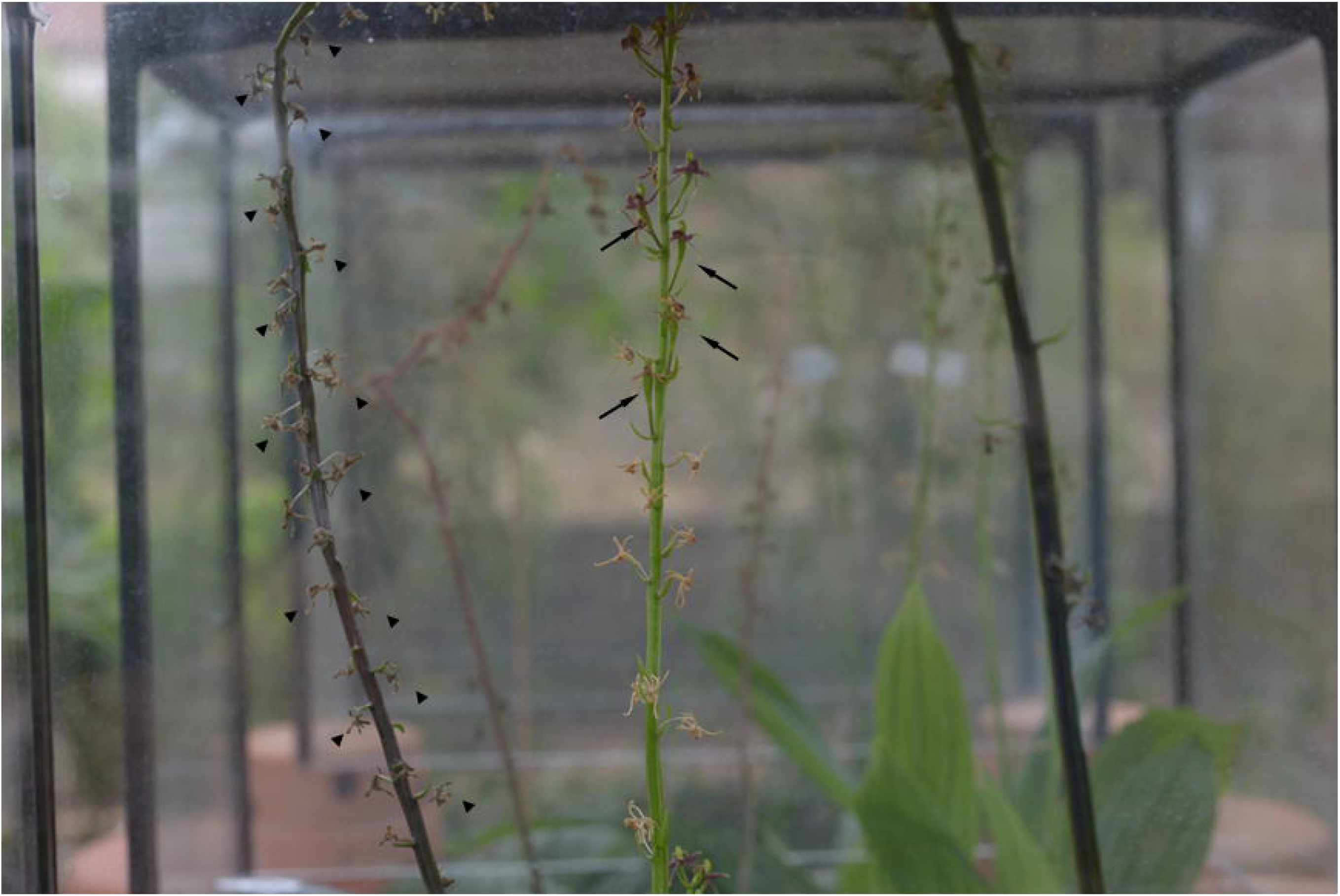
Details of the plants of *Liparis nervosa* and *Liparis* sp. inside the glass boxes (experiment of spontaneous self-pollination), where both rainfall and pollinators are excluded. Note the fruits of *Liparis* sp. (arrows), whereas in the inflorescences of *L. nervosa* no fruit is recorded (arrowheads).

### Phylogenetic analyses

The tree statistics for both combined and individual data sets from MP analyses, including characters in data matrices, scores of retention and consistency indices, numbers of characters, tree lengths, and most parsimonious trees are given in Appendix S3.

MP analyses of ITS and *matK-trnK* regions yielded 376 and 238 phylogenetically informative characters, respectively. Numbers of most parsimonious trees were 28 for ITS, and 669 for *matK-trnK* (Appendix S3). Phylogenetic analysis combining sequences of ITS (nrDNA) and *matK-trnK* (cpDNA) regions resulted in 18 most parsimonious trees, with 593 informative characters (Appendix S3).

### Maximum parsimony analysis of ITS (nrDNA)

The MP analysis obtained from isolated ITS matrix yielded a relatively well resolved strict consensus tree, but with a large polytomy embracing the *Liparis* with habit predominantly epiphytic plus *Oberonia* (BS 97; Appendix S4). *Dienia* emerges within the clade including the terrestrial *Liparis* with flat leaves (BS 99). The terrestrial species of *Liparis* with plicate leaves plus *Malaxis* and *Crepidium* form a well-supported clade (BS 100). The Asiatic specimen of *L. nervosa* emerges as sister to a large clade that includes the Brazilian *Liparis, Crepidium* and *Malaxis* with low support (BS 63, Appendix S4). The unidentified marshy species of *Liparis* emerges as sister to the Brazilian *L. nervosa* with strong support (BS 100; Appendix S4).

### Maximum parsimony analysis of *matK-trnK* (cpDNA)

As in the ITS tree, the isolated *matK-trnK* matrix yielded a relatively well resolved strict consensus tree, but with a large polytomy embracing the *Liparis* with habit predominantly epiphytic plus *Oberonia* (BS 73; Appendix S5). Terrestrial *Liparis* with plicate leaves form a well-supported clade that includes *Malaxis* and *Crepidium* (BS 100). *Dienia* emerges within the clade including the terrestrial *Liparis* with flat leaves (BS 100). The unidentified marshy species of *Liparis* emerges as sister to the Brazilian *L. nervosa* (BS 100; Appendix S5). The Brazilian clade *L. nervosa/L*. sp. is sister to the clade that includes the Asian *Liparis* with plicate leaves and terrestrial habit, *Crepidium* and *Malaxis* with low support (BS 52, Appendix S5). In the isolated *mat-K* analysis the Asiatic specimen of *L. nervosa* emerges in a clade that includes *L. layardii* and *L. formosana* (BS 96; Appendix S5).

### Phylogenetic analyses combining the regions ITS (nrDNA) and *matK-trnK* (cpDNA)

The trees of the analyses combining both regions are almost completely resolved. As in the isolated analyses, in all combined MP (Appendix S6), BI and ML analyses (Figs. 6–7), *Liparis* emerges as a polyphyletic genus. *Dienia* emerges within the clade including the terrestrial *Liparis* with flat leaves. As in all isolated analyses the terrestrial species of *Liparis* with plicate leaves, *Malaxis* and *Crepidium* form a well-supported clade (Appendix S6; Figs. 6–7). The remaining *Liparis* form a polytomy within a clade including *Oberonia*. *Liparis nervosa* emerges within a clade containing the terrestrial species with plicate leaves (Appendix S6; Figs. 6–7). The Asiatic *L. nervosa* is nested as sister to the clade embracing the Brazilian *L. nervosa* and *Liparis* sp. in MP, ML and BI analyses (BS 100, PP 1; Fig. 7). The unidentified marshy species of *Liparis* emerges as sister to the Brazilian specimens of *L. nervosa* with strong support (BS 100, PP 1; Figs. 6–7).

**FIGURE 5.**
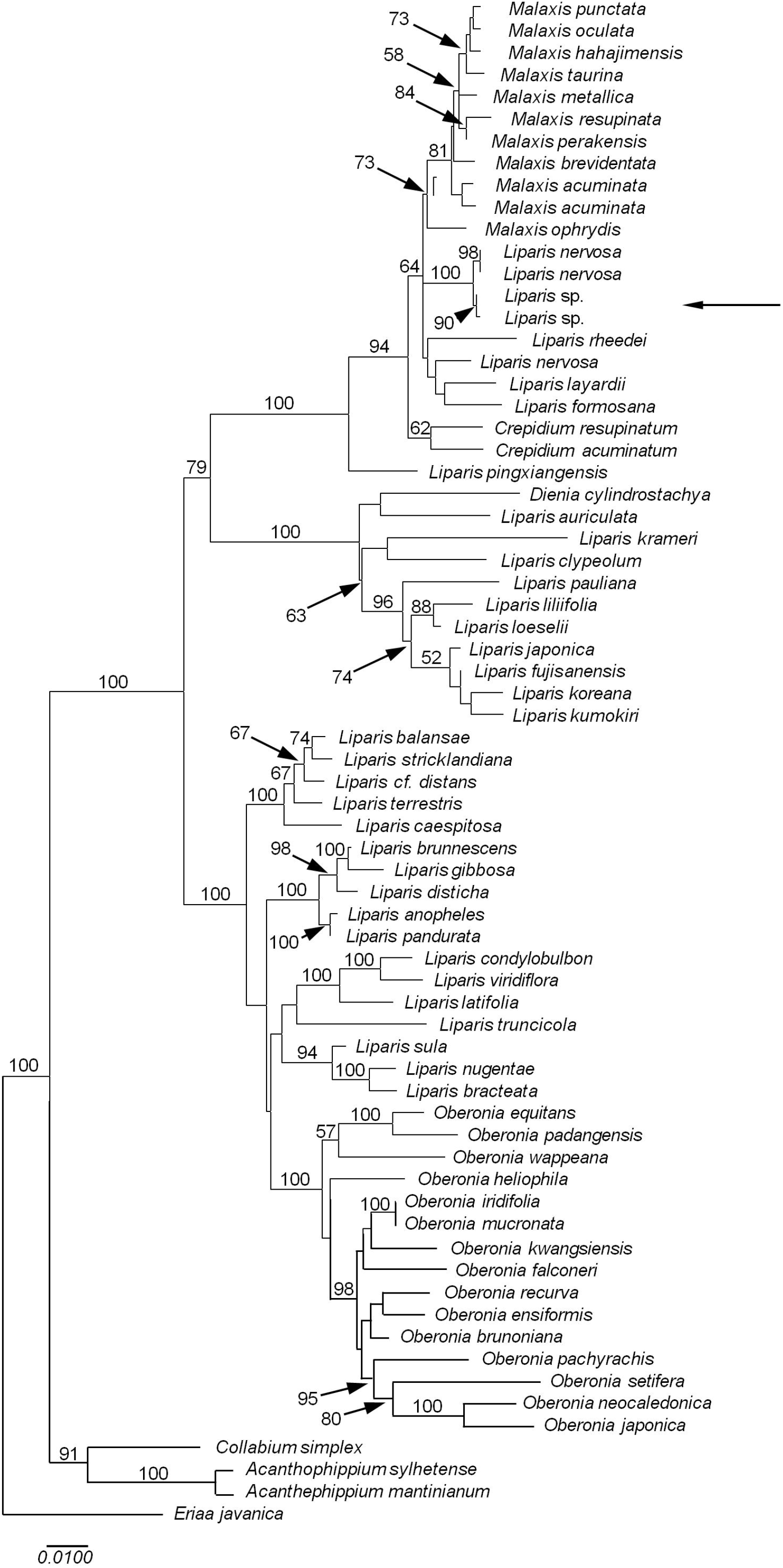
Maximum Likelihood analysis of *Liparis* Rich. (Orchidaceae) and outgroups based on the combination of the regions ITS (nrDNA) and *matK-trnK* (cpDNA). The tree with the highest log likelihood (−9757.5629) is shown. Initial tree(s) for the heuristic search was obtained automatically by applying Heuristic Search algorithm to a matrix of pairwise distances estimated using the Maximum Composite Likelihood (MCL) approach, and then selecting the topology with superior log likelihood value. The tree is drawn to scale, with branch lengths measured in the number of substitutions per site. Bootstrap/parsimony values >50 are given on branches. The position of *Liparis* sp. is given within plicate-leaved *Liparis* (arrow).

**FIGURE 6.**
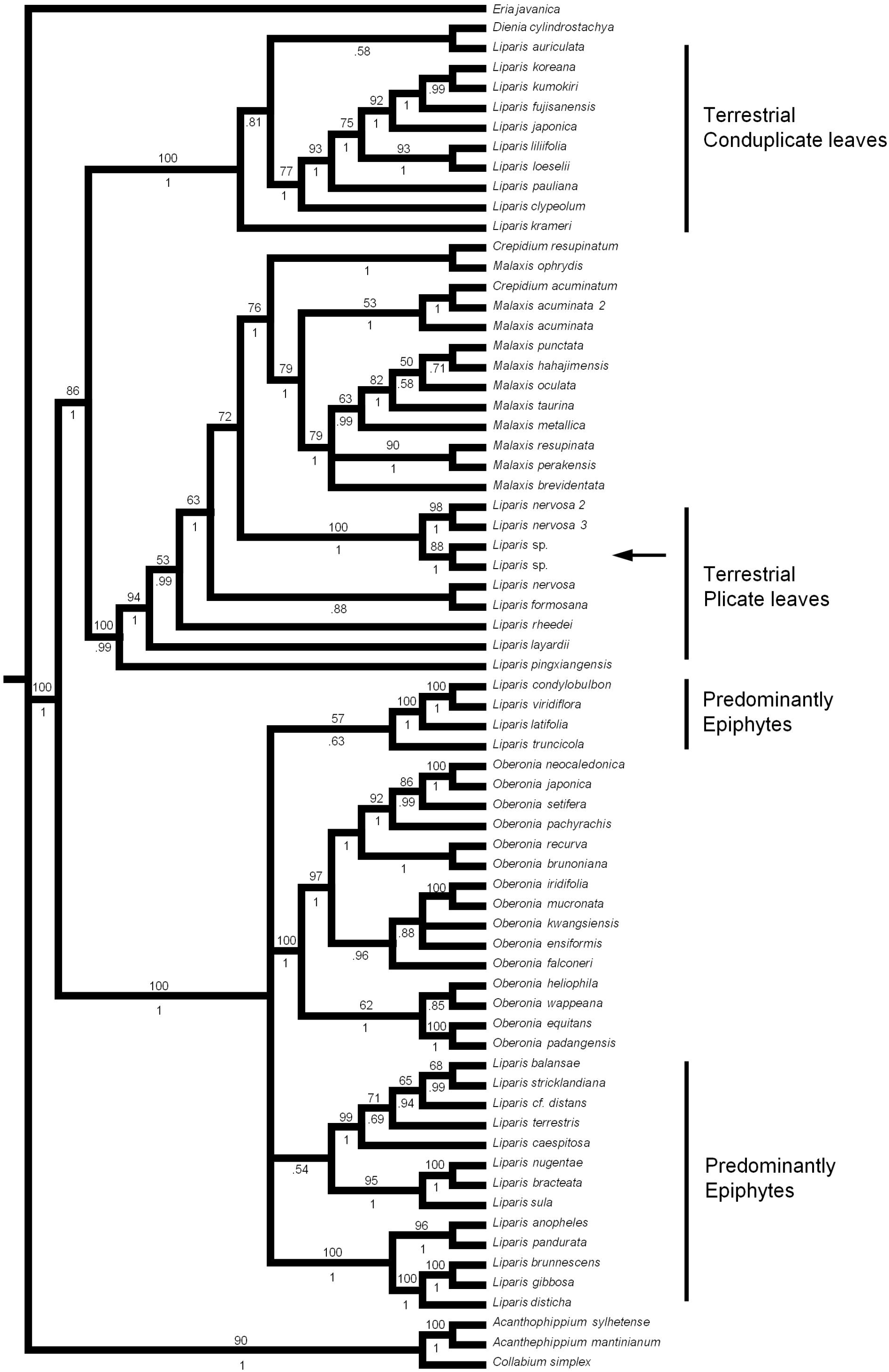
Bayesian inference analysis of *Liparis* (Orchidaceae) and outgroups based on combined ITS (nrDNA), and *matK-trnK* (cpDNA). Posterior probabilities values > 0.5 (BI) are given below branches. Bootstrap values >50 obtained from the MP analysis (matrix based on the combination of both ITS and *matK-trnK* regions) are given above branches. Note *Liparis* emerges as a polyphyletic genus, and the Brazilian *Liparis* sp. is included in the clade including terrestrial *Liparis* with plicate leaves (arrow).

**FIGURE 7.**
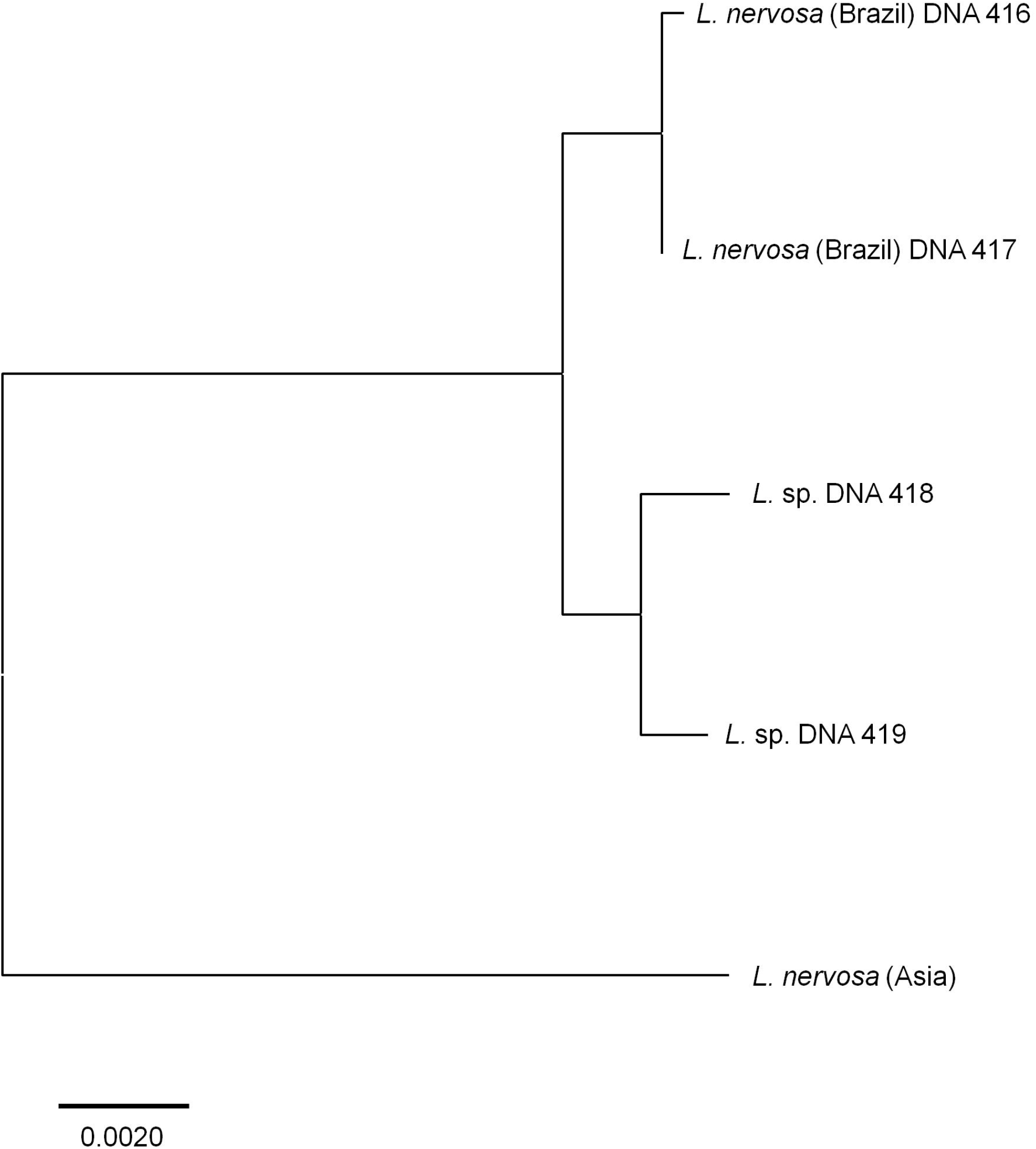
Molecular distances between *Liparis nervosa* and *Liparis* sp. based on usual plant DNA barcoding. The evolutionary history was inferred using the Neighbor-Joining method. The optimal tree with the sum of branch length = 0.02295621 is shown. The tree is drawn to scale, with branch lengths in the same units as those of the evolutionary distances used to infer the phylogenetic tree. The evolutionary distances were computed using the Kimura 2-parameter method and are in the units of the number of base substitutions per site.

### DNA barcoding analysis

The evolutionary distances between specimens of *Liparis nervosa* and *Liparis* sp., inferred by using the Neighbor-Joining method, resulted in an optimal tree with the sum of branch length = 0.02295621 (Appendix S7). In this tree, very similar ranges of intrapopulation and intra-specific sequence divergence were found for the combined cpDNA and nrDNA data matrix (Fig. 5). The estimates of distance divergence between sequences of specimens of an Asian *Liparis nervosa* and plants of *L. nervosa* and *Liparis* sp. collected in the study area (Jundiaí, southeastern Brazil), are presented in Appendix S7.

## DISCUSSION

Generic circumscription within Malaxidae is still unclear, since *Liparis* is polyphyletic and includes three groups (Cameron, 2005; Tsutsumi et al., 2007; Li and Yan, 2013): one clade with linear to lanceolate and conduplicate leaves, and a predominantly epiphyte emerging as sister to *Oberonia;* a second clade with terrestrial habit and usually 1-2 conduplicate leaves sheeting the inflorescence scape, related to *Dienia*; and a third group characterized by its terrestrial habit and plicate leaves, closely related to *Crepidium* and *Malaxis. Liparis* sp. emerges within plicate-leaved *Liparis* and sister to the Brazilian *L. nervosa*, whereas the Asiatic *L. nervosa* is sister to the Brazilian clade including *L. nervosa/L*. sp. with strong support (BS 100, PP 1). Although we have no access to Asiatic populations of *L. nervosa*, a detailed study on morphology and experimental taxonomy can be important along their distribution, since our analysis suggests the Brazilian *L. nervosa* is a distinct taxon, possibly *L. elata* Lindl. Currently, *Liparis nervosa* is been considered as a widespread species, occurring throughout tropical Asia, Africa and America, and the type specimen was collected near Osaka, Japan (Lindley, 1830).

Our study shows *L. nervosa* is dependent on a biotic pollen vector for pollination, while the flowers of sympatric congener *Liparis* sp. self-pollinates. The emergence of autogamous lineages derivate from pollinator-dependent species is a common situation in orchids with dispersion from the continent to islands. Since the orchid seeds are capable of dispersion over great distances by wind, species can be established in islands where pollination service can be chronically deficient or absent. In fact, some of these species are xenogamous on the continent and autogamous on the islands (Aguiar et al., 2012). Due the deficient pollination service on the island, selection occurs in plants able to self-pollination. The divergence from parental population can result in a future speciation event. This is a good model to explain allopatric speciation. In fact, the existence of prezygotic barriers is the main factor impeding the gene flow between distinct taxa. When pre-mating barriers are broken or do not exist, gene flow between distinct species can occurs and hybrid progenies can arise (Pansarin and Amaral, 2008). When hybrid progeny is locally adapted and reproductive isolate from parental due pre- or post-mating barriers at a later time a speciation event can occurs. In fact, natural hybridization is the simplest pathway for sympatric speciation. However, how we demonstrate in this paper, species can arisen can arise in simpatry without a hybrid origin. This is the most plausible hypothesis, since the unique species of *Liparis* occurring in the Brazilian forests is *Liparis nervosa* (Barros et al., 2019). In addition, the paludal *Liparis* shows vegetative and floral morphology closely related to *L. nervosa*. Besides these evidences, other factor suggests the paludal *Liparis* is a sympatric cryptic species: (1) *Liparis nervosa* and *Liparis* sp. are currently sympatric; (2) both sympatric species are isolated reproductively due divergence in flowering period, occurrence in distinct microhabitats and reproductive strategies; (3) the allopatric isolation is actually improbable; (4) they are monophyletic and represented as sister taxa in the phylogenies; and (5) are genetically distinct taxa (Les et al., 2015).

Although flower characteristics have been extensively used in plant taxonomy, including Orchidaceae (Dressler, 1993; Pridgeon et al., 1999), groups of Malaxidae can be easily recognized on the basis of their vegetative characters (Cameron, 2005; Li and Yan, 2013). Conversely, flower morphology is little informative, showing strong similarity concerning shape and colour. Within *Liparis*, species belonging to distinct clades are practically indistinguishable with regard to morphology of flowers. In fact, Brazilian *Liparis* are easily recognized when their vegetative characters are compared, but the morphology of their flowers is similar. This is not uncommon, since many orchid genera show strong similarity in flowers characteristics among the species (e.g. *Stelis* Sw., *Polystachya* Hook., *Cleistes* Rich.). Furthermore, parallelisms in flower characteristics involving distinct genera within Orchidaceae and between orchids and other plant families frequently occur (Vale et al., 2011; Pansarin et al., 2017).

While sympatric speciation seems to be common in orchids, species are reproductively isolated due differences in flower morphology and olfactory plus color signals (Xu et al., 2012). These differences are easily recognized by usual methods in plant taxonomy. Conversely, in cryptic sympatric speciation, often these morphological markers are obscure and the species distinction based exclusively on plant phenotypes can be difficult. Nevertheless, how to recognize a cryptic sympatric species? As demonstrated here for *Liparis* sp., the combination of molecular approach plus detailed investigation on plant morphology based on living plants, study on microhabitats, flowering phenology and breeding system can elucidate the identity of a cryptic species. A cryptic plant species can also be distinguishable based on study about its natural history, including pollinators, pollination mechanisms, breeding systems and morpho-anatomy of flowers (Pansarin and Amaral, 2006). Indeed, our study demonstrates the importance of multiple data sources to resolve the status of a potential cryptic plant species.

Of course, the decisions on species boundaries are dependent on which species concept is being used (Tan et al., 2010), since at least 22 species concepts have been listed (Mayden, 1997). However, these species concepts can be grouped in four main categories (Tan et al., 2010). One of the four categories is the Phylogenetic Species Concept, which is based on monophyletic groups (Mishler and Theriot, 2000). The second category is the Biological Species Concept, which is based on reproductive isolation between organisms (Mayr, 2000), and the Hennigian Species Concept (Meier and Willmann, 2000). The remaining two categories are the concepts with group the concepts based on the recognition of populations, i.e. Phylogenetic Species Concept (Wheeler and Platnick, 2000), and the category that use a combination of criterion, namely Evolutionary Species Concept (Wiley and Mayden, 2000).

Since reproductive isolation is the principle for Hennigian plus Biological Species Concepts, our data suggest the cryptic *Liparis* sp. and *L. nervosa* are reproductive isolate due their distinct flowering seasons, microhabitats and reproduction strategies, although both are sympatric. Furthermore, our data show the cryptic *Liparis* sp. and *L. nervosa* can be considered as distinct taxa based on DNA barcoding analysis and in the phylogenetic approach based on the combination of nrDNA and cpDNA regions, suggesting both species can be separate each other on basis on both Phylogenetic Species Concept (Wheeler and Platnick, 2000), and the Evolutionary Species Concept (Wiley and Mayden, 2000). Although plant DNA barcoding is an important tool in the recognizing species boundaries, these clusters are more adequate to the Phylogenetic Species Concept and are not designed to the Biological Species Concept. Furthermore, when the circularity in the alpha taxonomy is signalized by the barcoding approach, supplementary methods, such as experimental taxonomy, are necessary to draw concrete conclusions about species identity (Chen et al., 2011).

## CONCLUSIONS

Although formerly recognized as a unique species, *Liparis nervosa* comprised two distinct sister taxa. According to our data, we conclude the paludal *Liparis* is a cryptic sympatric taxon, distinguishable from *L. nervosa* by differences in habitat, flowering period, vegetative plus floral morphology (recognized when analyses are based on fresh material), and reproduction strategies. These evidences are supported by molecular investigations. *Liparis* sp. emerges as sister to *L. nervosa*, within a clade of species with terrestrial habitat and plicate leaves. The discovery of sympatric cryptic speciation in orchids provides increase on the knowledge of diversification and reproductive isolation in plants.

## ACKNOWLEDGEMENTS

The authors thank the Base Ecológica da Serra do Japi and Guarda Municipal de Jundiaí for permission for the fieldwork. Provision of funds by FAPESP (grant 14/14969-6 and grant 18/07357-5).

## AUTHORS’ CONTRIBUTIONS

ERP developed the idea of the study, performed the experiments and wrote the manuscript; AWCF participated in describing the results. All authors read and approved the final version of the manuscript.

## SUPPORTING INFORMATION

**APPENDIX S1.** Species of *Liparis* (Orchidaceae) and outgroups included in the molecular studies, vouchers, and GenBank accession numbers for the regions ITS (nrDNA), *matK-trnK* (cpDNA).

**APPENDIX S2.** Effect of air temperature and relative humidity on occurrence of spontaneous self-pollination on flowers of *Liparis nervosa* and *Liparis* sp. maintained under glass boxes (flowers protected from water and pollinators).

**APPENDIX S3.** Summary of results of maximum parsimony analyses of *Liparis* (Orchidaceae, Epidendroideae) and outgroups

**APPENDIX S4.** Strict consensus tree based on maximum parsimony analysis from isolated ITS (nrDNA) sequences of *Liparis* (Orchidaceae) and outgroups. Bootstrap values >50 are given on branches.

**APPENDIX S5.** Strict consensus tree based on maximum parsimony analysis from isolated *matK-trnK* (cpDNA) sequences of *Liparis* (Orchidaceae) and outgroups. Bootstrap values >50 are given on branches.

**APPENDIX S6.** Strict consensus tree based on maximum parsimony analysis from combined ITS and *matK-trnK* sequences of *Liparis* (Orchidaceae) and outgroups. Bootstrap values >50 are given on branches.

**APPENDIX S7.** Evolutionary distances between specimens of *Liparis nervosa* and *Liparis* sp., inferred by using the Neighbor-Joining method and Kimura 2-parameter model.

