## Supplementary material for "Elucidating cryptic sympatric speciation in terrestrial orchids": S1

**Appendix 1**. Species of *Liparis* (Orchidaceae) and outgroups included in the molecular studies, vouchers, and GenBank accession numbers for the regions ITS (nrDNA), *matK-trnK* (cpDNA).

| Species | Voucher number | ITS | *matK* |
| --- | --- | --- | --- |
| *Acanthephippium mantinianum* L.Linden & Cogn |  | AF521081* | AF263618* |
| *Acanthophippium sylhetense* Lindl. | Z.J. Liu 3790 | KM025144* | KF673783* |
| *Collabium simplex* Rchb.f. | AK991017/1/04 | EF670387* |  |
| *Collabium simplex* Rchb.f. |  |  | AY557200* |
| *Crepidium acuminatum* (D. Don) Szlach. | SBB-0890 | JN114482* |  |
| *Crepidium acuminatum* (D. Don) Szlach. | Z.J.Liu 5425 |  | KJ459304* |
| *Crepidium resupinatum* (G. Forst.) Szlach. | SBB-0952 |  | JN004410* |
| *Crepidium resupinatum* (G. Forst.) Szlach. |  | KM583453* |  |
| *Dienia cylindrostachya* Lindl. | BB-0907 | JN114492* | JN004423* |
| *Eria javanica* (Sw.) Blume | CBG 740854 | AY240014* |  |
| *Eria javanica* (Sw.) Blume | Szlachetko s.n. |  | EF079354* |
| *Liparis anopheles* J.J. Wood | Leiden cult. 980165 | AY907075* | AY907139* |
| *Liparis auriculata* Blume ex Miq. | TBG144267 (TNS) | AB289458* |  |
| *Liparis auriculata* Blume ex Miq. |  |  | KF262076* |
| *Liparis balansae* Gagnep. | L. Li 152 | KF589874* | KF589880* |
| *Liparis bracteata* D.G. Hunt | P. Weston s.n. | AY907076* | AY907140* |
| *Liparis brunnescens* Schltr. | Leiden cult. 20030224 | AY907098* | AY907165* |
| *Liparis caespitosa* (Thouars) Lindl. | Leiden cult. 20030195 | AY907077* | AY907141* |
| *Liparis clypeolum* (G. Forst.) Lindl. | J.-Y. Meyer 1029 (NY) | AY907079* | AY907143* |
| *Liparis condylobulbon* Rchb. f. | Leiden cult. 20030654 | AY907080* | AY907144* |
| *Liparis cf. distans* C.B. Clarke |  | AJ558105* |  |
| *Liparis distans* C.B. Clarke | ELipD |  | KF361627* |
| *Liparis disticha* (Thouars) Lindl. | Leiden cult. 20010180 | AY907081* | AY907145* |
| *Liparis formosana* Rchb. f. | K. Cameron 2151 (NY) | AY907082* | AY907147* |
| *Liparis fujisanensis* Maek. | Lee 239 (EWH) | EU024936* |  |
| *Liparis fujisanensis* Maek. | Tsutsumi L18 (TNS) |  | AB289511* |
| *Liparis gibbosa* Finet | K. Cameron 2211 (NY) | AY907084* | AY907149* |
| *Liparis japonica* Maxim. | K. Cameron 2176 (NY) | AY907086* | AY907151* |
| *Liparis koreana* (Nakai) Nakai | EWH:Lee 200 | EU017423* |  |
| *Liparis koreana* (Nakai) Nakai | Tsutsumi L4 (TNS) |  | AB289515* |
| *Liparis krameri* Franch. & Sav. | Tsutsumi L21(TNS) | AB289469* |  |
| *Liparis krameri* Franch. & Sav. |  |  | KC704647* |
| *Liparis kumokiri* F. Maek. | K. Cameron 2147 (NY) | AY907087* | AY907152* |
| *Liparis latifolia* Lindl. | Singapore B. G. cult. 837 | AY907088* | AY907153* |
| *Liparis layardii* F.Muell. | K. Cameron 2060 (NY) | AY907089* | AY907155* |
| *Liparis liliifolia* (L.) Rich. ex Ker Gawl. | C. McCartney s.n. | AY907090* |  |
| *Liparis liliifolia* (L.) Rich. ex Ker Gawl. | Tsutsumi L25 (TNS) |  | AB289521* |
| *Liparis loeselii* (L.) Rich. | B. Ewacha s.n. | AY907091* | AY907157* |
| *Liparis nervosa* (Thunb.) Lindl. | K. Cameron 2150 (NY) | AY907092* | AY907158* |
| *Liparis nervosa* (Thunb.) Lindl. | SBB-0071 |  |  |
| *Liparis nervosa* (Thunb.) Lindl. |  |  |  |
| *Liparis nervosa* (Thunb.) Lindl. | E. R. Pansarin s.n. DNA 416 (LBMBP) |  |  |
| *Liparis nervosa* (Thunb.) Lindl. | E. R. Pansarin s.n. DNA 417 (LBMBP) |  |  |
| *Liparis nugentae* F.M. Bailey | P. Weston s.n. | AY907093* | AY907159* |
| *Liparis pandurata* Ames | Singapore B. G. cult. 85 | AY907095* | AY907161* |
| *Liparis pauliana* Hand.-Mazz | K. Cameron 2169 (NY) | AY907096* | AY907163* |
| *Liparis pingxiangensis* L. Li & H. F. Yan | L. Li 157 | KF589872* | KF589878* |
| *Liparis rheedii* (Blume) Lindl. | Leiden cult. 970454 | AY907097* | AY907164* |
| *Liparis* sp. “cryptic” | E. R. Pansarin s.n. DNA 418 (LBMBP) |  |  |
| *Liparis* sp.“cryptic” | E. R. Pansarin s.n. DNA 419 (LBMBP) |  |  |
| *Liparis sula* N. Hallé | K. Cameron s.n. DNA#1174 | AY907104* | AY907171* |
| *Liparis stricklandiana* Rchb. f. | Z.J.Liu 4341 | KJ459298* | KJ459328* |
| *Liparis terrestris* J.B. Comber | Singapore B. G. cult. 3482 | AY907105* | AY907172* |
| *Liparis truncicola* Schltr. | Leiden cult. 20030222 | AY907106* | AY907173* |
| *Liparis viridiflora* [(Blume) Lindl.](http://www.tropicos.org/NamePage.aspx?nameid=23500829) | NYBG cult. 2025 | AY907107* | AY907174* |
| *Malaxis acuminata* D. Don | M. Watanabe s. n. (TNS) | AB290884* | AB290892* |
| *Malaxis acuminata* D. Don | SBB-1564 | KX277725* | KX344572* |
| *Malaxis brevidentata* C. Schweinf. | Yukawa 97-2066 (TNS) | AB290886* | AB290894* |
| *Malaxis hahajimensis* S. Kobay. | Ohi-Toma s. n. (TI) | AB290888* | AB290896* |
| *Malaxis metallica* (Rchb. f.) Kuntze | Singapore B. G. cult. 1522 | AY907113* | AY907180* |
| *Malaxis oculata* (Rchb. f.) Kuntze | Nazarudin s. n. (TNS) | AB290890* | AB290898* |
| *Malaxis ophrydis* (J. Koenig) Ormerod | Singapore B. G. cult. 3380 | AY907114* | AY907181* |
| *Malaxis perakensis* (Ridl.) Holttum | Nazarudin s. n. (TNS) | AB290891* | AB290899* |
| *Malaxis punctata* J.J. Wood | Leiden cult. 980113 | AY907117* | AY907184* |
| *Malaxis resupinata* Kuntze | K. Wood 9548 (NY) | AY907118* | AY907185* |
| *Malaxis taurina* (Rchb. f.) Kuntze | T. Motley & K. Cameron 2150 (NY) | AY907128* | AY907192* |
| *Oberonia brunoniana* Wight | SBB-0795 | JN114625* | JN004519* |
| *Oberonia ensiformis* (Sm.) Lindl. | SBB-0798 | JN114628* | JN004522* |
| *Oberonia equitans* (Thouars) Schltr. | T. Motley & K. Cameron 2255 (NY) | AY907130* | AY907198* |
| *Oberonia falconeri* Hook. f. | SBB-0794 | JN114633* | JN004527* |
| *Oberonia heliophila* Rchb. f. | T. Motley & K. Cameron 2243 (NY) | AY907131* | AY907199* |
| *Oberonia iridifolia* Lindl. | NYBG cult. 107/67 | AY907132* | AY907200* |
| *Oberonia japonica* (Maxim.) Makino | Z.J. Liu 5216 | KF560543* | KF673836* |
| *Oberonia kwangsiensis* Seidenf. | Z.J. Liu 4082 | KF560542* | KF673837* |
| *Oberonia mucronata* (D. Don) Ormerod & Seidenf. | SBB-0234 | JN114641* | JN004535* |
| *Oberonia neocaledonica* Schltr. | T. Motley & K. Cameron 2173 (NY) | AY907134* | AY907202* |
| *Oberonia pachyrachis* Rchb. f. ex Hook. f. | SBB-0265 | JN114645* | JN004538* |
| *Oberonia padangensis* Schltr. | Leiden cult. 960231 | AY907135* | AY907203* |
| *Oberonia setifera* Lindl. | NYBG cult. s.n. ex Andy's Orchids | AY907136* | AY907204* |
| *Oberonia recurva* Lindl. | SBB-0800 | JN114648* |  |
| *Oberonia recurva* Lindl. | SBB-0801 |  | JN004544* |
| *Oberonia wappeana* J.J. Sm. | Leiden cult. 20030243 | AY907138* | AY907206* |

* Sequences taken from Genbank.
