## Supplementary material for "Elucidating cryptic sympatric speciation in terrestrial orchids": S2

**Appendix 2**. Effect of air temperature and relative humidity on occurrence of spontaneous self-pollination on flowers of *Liparis nervosa* and *Liparis* sp. maintained under glass boxes (flowers protected from water and pollinators).

Relative humidity %

Temperature °C

| From 13 November 2015 to 15 February 2016 | RELATIVE-HUMIDITY |
| --- | --- |
| TIME | 48.2 |
| 15:38:11 | 57.7 |
| 16:38:11 | 60.6 |
| 17:38:11 | 64.1 |
| 18:38:11 | 74.7 |
| 19:38:11 | 82 |
| 20:38:11 | 72.7 |
| 21:38:11 | 70 |
| 22:38:11 | 70.4 |
| 23:38:11 | 74.6 |
| 00:38:11 | 78.2 |
| 01:38:11 | 82.8 |
| 02:38:11 | 87 |
| 03:38:11 | 89.7 |
| 04:38:11 | 91.4 |
| 05:38:11 | 89.5 |
| 06:38:11 | 84.6 |
| 07:38:11 | 76.9 |
| 08:38:11 | 56.4 |
| 09:38:11 | 53.2 |
| 10:38:11 | 50.6 |
| 11:38:11 | 49.9 |
| 12:38:11 | 47.8 |
| 13:38:11 | 46.8 |
| 14:38:11 | 49.6 |
| 15:38:11 | 72.6 |
| 16:38:11 | 64.7 |
| 17:38:11 | 62.7 |
| 18:38:11 | 73.4 |
| 19:38:11 | 78.4 |
| 20:38:11 | 77.6 |
| 21:38:11 | 79.7 |
| 22:38:11 | 82.5 |
| 23:38:11 | 79.5 |
| 00:38:11 | 78.2 |
| 01:38:11 | 88.6 |
| 02:38:11 | 92.2 |
| 03:38:11 | 93.3 |
| 04:38:11 | 93.6 |
| 05:38:11 | 94.7 |
| 06:38:11 | 94.7 |
| 07:38:11 | 92.7 |
| 08:38:11 | 82.2 |
| 09:38:11 | 75.2 |
| 10:38:11 | 67.7 |
| 11:38:11 | 62.8 |
| 12:38:11 | 64.2 |
| 13:38:11 | 57.8 |
| 14:38:11 | 59 |
| 15:38:11 | 58.9 |
| 16:38:11 | 45.6 |
| 17:38:11 | 70.5 |
| 18:38:11 | 84.7 |
| 19:38:11 | 89.8 |
| 20:38:11 | 91.4 |
| 21:38:11 | 92.4 |
| 22:38:11 | 93.6 |
| 23:38:11 | 94.1 |
| 00:38:11 | 94.1 |
| 01:38:11 | 94.5 |
| 02:38:11 | 94.6 |
| 03:38:11 | 94.7 |
| 04:38:11 | 95 |
| 05:38:11 | 94.8 |
| 06:38:11 | 86.6 |
| 07:38:11 | 75.5 |
| 08:38:11 | 59.6 |
| 09:38:11 | 57.7 |
| 10:38:11 | 58.6 |
| 11:38:11 | 78.2 |
| 12:38:11 | 78 |
| 13:38:11 | 96.6 |
| 14:38:11 | 98 |
| 15:38:11 | 98.4 |
| 16:38:11 | 97.6 |
| 17:38:11 | 98.1 |
| 18:38:11 | 98.3 |
| 19:38:11 | 98.5 |
| 20:38:11 | 98.6 |
| 21:38:11 | 98.7 |
| 22:38:11 | 98.6 |
| 23:38:11 | 98.8 |
| 00:38:11 | 98.8 |
| 01:38:11 | 98.8 |
| 02:38:11 | 98.7 |
| 03:38:11 | 98.8 |
| 04:38:11 | 98.9 |
| 05:38:11 | 99.2 |
| 06:38:11 | 95.7 |
| 07:38:11 | 89.9 |
| 08:38:11 | 83.4 |
| 09:38:11 | 74 |
| 10:38:11 | 67.9 |
| 11:38:11 | 76 |
| 12:38:11 | 60 |
| 13:38:11 | 58.1 |
| 14:38:11 | 77.2 |
| 15:38:11 | 79.5 |
| 16:38:11 | 72.9 |
| 17:38:11 | 83.4 |
| 18:38:11 | 79.4 |
| 19:38:11 | 84.4 |
| 20:38:11 | 84.7 |
| 21:38:11 | 89.4 |
| 22:38:11 | 92.2 |
| 23:38:11 | 94.3 |
| 00:38:11 | 95.1 |
| 01:38:11 | 95.8 |
| 02:38:11 | 96.5 |
| 03:38:11 | 96.6 |
| 04:38:11 | 96.7 |
| 05:38:11 | 96.9 |
| 06:38:11 | 96.1 |
| 07:38:11 | 81.2 |
| 08:38:11 | 63.6 |
| 09:38:11 | 60.3 |
| 10:38:11 | 51.3 |
| 11:38:11 | 55 |
| 12:38:11 | 49.9 |
| 13:38:11 | 48.7 |
| 14:38:11 | 50.5 |
| 15:38:11 | 55.1 |
| 16:38:11 | 53.9 |
| 17:38:11 | 67.2 |
| 18:38:11 | 72.8 |
| 19:38:11 | 77.6 |
| 20:38:11 | 80.8 |
| 21:38:11 | 81.2 |
| 22:38:11 | 84.6 |
| 23:38:11 | 88.4 |
| 00:38:11 | 89.6 |
| 01:38:11 | 88.6 |
| 02:38:11 | 89.4 |
| 03:38:11 | 91 |
| 04:38:11 | 91.6 |
| 05:38:11 | 90.8 |
| 06:38:11 | 84.6 |
| 07:38:11 | 79.3 |
| 08:38:11 | 61.7 |
| 09:38:11 | 63.7 |
| 10:38:11 | 56.1 |
| 11:38:11 | 52.1 |
| 12:38:11 | 51.7 |
| 13:38:11 | 77.5 |
| 14:38:11 | 69.7 |
| 15:38:11 | 76.3 |
| 16:38:11 | 63.2 |
| 17:38:11 | 78.4 |
| 18:38:11 | 84.2 |
| 19:38:11 | 88.4 |
| 20:38:11 | 91.5 |
| 21:38:11 | 92.1 |
| 22:38:11 | 93.7 |
| 23:38:11 | 94.5 |
| 00:38:11 | 94.5 |
| 01:38:11 | 94.1 |
| 02:38:11 | 93.9 |
| 03:38:11 | 93.3 |
| 04:38:11 | 92.9 |
| 05:38:11 | 96.6 |
| 06:38:11 | 95.2 |
| 07:38:11 | 96.8 |
| 08:38:11 | 97.5 |
| 09:38:11 | 96.2 |
| 10:38:11 | 93.2 |
| 11:38:11 | 91 |
| 12:38:11 | 83.8 |
| 13:38:11 | 81.7 |
| 14:38:11 | 81.8 |
| 15:38:11 | 81.7 |
| 16:38:11 | 80.5 |
| 17:38:11 | 86.9 |
| 18:38:11 | 91.2 |
| 19:38:11 | 93.1 |
| 20:38:11 | 94.8 |
| 21:38:11 | 95.9 |
| 22:38:11 | 96.6 |
| 23:38:11 | 97.3 |
| 00:38:11 | 97.5 |
| 01:38:11 | 97.9 |
| 02:38:11 | 98.3 |
| 03:38:11 | 98.6 |
| 04:38:11 | 98.7 |
| 05:38:11 | 99 |
| 06:38:11 | 98.4 |
| 07:38:11 | 91 |
| 08:38:11 | 72.1 |
| 09:38:11 | 70.7 |
| 10:38:11 | 61.5 |
| 11:38:11 | 58.6 |
| 12:38:11 | 52.1 |
| 13:38:11 | 52 |
| 14:38:11 | 54.7 |
| 15:38:11 | 71.9 |
| 16:38:11 | 58.6 |
| 17:38:11 | 74.3 |
| 18:38:11 | 84.1 |
| 19:38:11 | 88.5 |
| 20:38:11 | 90.8 |
| 21:38:11 | 92.6 |

|  | TEMPERATURE |
| --- | --- |
| TIME | 34.7 |
| 15:38:11 | 34.3 |
| 16:38:11 | 32.4 |
| 17:38:11 | 29.5 |
| 18:38:11 | 27.5 |
| 19:38:11 | 26 |
| 20:38:11 | 25.7 |
| 21:38:11 | 25.5 |
| 22:38:11 | 25 |
| 23:38:11 | 23.9 |
| 00:38:11 | 23.1 |
| 01:38:11 | 22.5 |
| 02:38:11 | 21.8 |
| 03:38:11 | 21.2 |
| 04:38:11 | 21 |
| 05:38:11 | 22.7 |
| 06:38:11 | 24.5 |
| 07:38:11 | 26.8 |
| 08:38:11 | 34 |
| 09:38:11 | 33.6 |
| 10:38:11 | 34.4 |
| 11:38:11 | 34.3 |
| 12:38:11 | 35.5 |
| 13:38:11 | 36 |
| 14:38:11 | 34.7 |
| 15:38:11 | 29.3 |
| 16:38:11 | 29.1 |
| 17:38:11 | 27.8 |
| 18:38:11 | 26.4 |
| 19:38:11 | 25.3 |
| 20:38:11 | 25.1 |
| 21:38:11 | 24.9 |
| 22:38:11 | 24.6 |
| 23:38:11 | 24.4 |
| 00:38:11 | 23.9 |
| 01:38:11 | 20.7 |
| 02:38:11 | 19.9 |
| 03:38:11 | 19.8 |
| 04:38:11 | 19.7 |
| 05:38:11 | 20 |
| 06:38:11 | 20.5 |
| 07:38:11 | 21.3 |
| 08:38:11 | 24.5 |
| 09:38:11 | 26.2 |
| 10:38:11 | 28.2 |
| 11:38:11 | 29.5 |
| 12:38:11 | 29.6 |
| 13:38:11 | 30.5 |
| 14:38:11 | 30.1 |
| 15:38:11 | 31.9 |
| 16:38:11 | 38.3 |
| 17:38:11 | 27.8 |
| 18:38:11 | 24.2 |
| 19:38:11 | 22.6 |
| 20:38:11 | 22 |
| 21:38:11 | 21.5 |
| 22:38:11 | 21.3 |
| 23:38:11 | 21.2 |
| 00:38:11 | 21.4 |
| 01:38:11 | 21.4 |
| 02:38:11 | 21.5 |
| 03:38:11 | 21.5 |
| 04:38:11 | 21.3 |
| 05:38:11 | 22.1 |
| 06:38:11 | 25 |
| 07:38:11 | 27.1 |
| 08:38:11 | 31.7 |
| 09:38:11 | 32.1 |
| 10:38:11 | 31 |
| 11:38:11 | 27.5 |
| 12:38:11 | 27.1 |
| 13:38:11 | 21 |
| 14:38:11 | 22.5 |
| 15:38:11 | 24.2 |
| 16:38:11 | 23.3 |
| 17:38:11 | 23 |
| 18:38:11 | 22.9 |
| 19:38:11 | 22.8 |
| 20:38:11 | 22.8 |
| 21:38:11 | 22.7 |
| 22:38:11 | 22.3 |
| 23:38:11 | 22.1 |
| 00:38:11 | 21.9 |
| 01:38:11 | 21.9 |
| 02:38:11 | 21.4 |
| 03:38:11 | 21 |
| 04:38:11 | 20.8 |
| 05:38:11 | 21.6 |
| 06:38:11 | 23.8 |
| 07:38:11 | 25.4 |
| 08:38:11 | 27.8 |
| 09:38:11 | 29.2 |
| 10:38:11 | 30.3 |
| 11:38:11 | 28.3 |
| 12:38:11 | 31 |
| 13:38:11 | 31.8 |
| 14:38:11 | 28 |
| 15:38:11 | 27.6 |
| 16:38:11 | 28.4 |
| 17:38:11 | 26.5 |
| 18:38:11 | 22.6 |
| 19:38:11 | 21 |
| 20:38:11 | 21.2 |
| 21:38:11 | 21.1 |
| 22:38:11 | 20.3 |
| 23:38:11 | 19.9 |
| 00:38:11 | 19.7 |
| 01:38:11 | 19.7 |
| 02:38:11 | 20.8 |
| 03:38:11 | 20.8 |
| 04:38:11 | 20.4 |
| 05:38:11 | 20.9 |
| 06:38:11 | 22.8 |
| 07:38:11 | 24.7 |
| 08:38:11 | 29.3 |
| 09:38:11 | 30.2 |
| 10:38:11 | 33.5 |
| 11:38:11 | 31 |
| 12:38:11 | 32.7 |
| 13:38:11 | 34.3 |
| 14:38:11 | 32.5 |
| 15:38:11 | 32 |
| 16:38:11 | 32.6 |
| 17:38:11 | 28.9 |
| 18:38:11 | 26.5 |
| 19:38:11 | 25.4 |
| 20:38:11 | 25 |
| 21:38:11 | 24.4 |
| 22:38:11 | 23.6 |
| 23:38:11 | 23 |
| 00:38:11 | 22.9 |
| 01:38:11 | 23.3 |
| 02:38:11 | 23 |
| 03:38:11 | 22.7 |
| 04:38:11 | 22.6 |
| 05:38:11 | 23.1 |
| 06:38:11 | 24.8 |
| 07:38:11 | 26.1 |
| 08:38:11 | 31.8 |
| 09:38:11 | 31.9 |
| 10:38:11 | 33.6 |
| 11:38:11 | 34 |
| 12:38:11 | 34.2 |
| 13:38:11 | 27.1 |
| 14:38:11 | 28.9 |
| 15:38:11 | 27.7 |
| 16:38:11 | 31.7 |
| 17:38:11 | 27.2 |
| 18:38:11 | 25.3 |
| 19:38:11 | 24.2 |
| 20:38:11 | 23.3 |
| 21:38:11 | 22.6 |
| 22:38:11 | 22.2 |
| 23:38:11 | 21.9 |
| 00:38:11 | 21.8 |
| 01:38:11 | 21.8 |
| 02:38:11 | 22.1 |
| 03:38:11 | 22.3 |
| 04:38:11 | 22.3 |
| 05:38:11 | 21.7 |
| 06:38:11 | 19.8 |
| 07:38:11 | 19.6 |
| 08:38:11 | 20.2 |
| 09:38:11 | 21.3 |
| 10:38:11 | 22.3 |
| 11:38:11 | 22.7 |
| 12:38:11 | 24.9 |
| 13:38:11 | 25.7 |
| 14:38:11 | 25.7 |
| 15:38:11 | 26 |
| 16:38:11 | 26.1 |
| 17:38:11 | 24.2 |
| 18:38:11 | 22.9 |
| 19:38:11 | 22.4 |
| 20:38:11 | 21.8 |
| 21:38:11 | 21.1 |
| 22:38:11 | 20.6 |
| 23:38:11 | 20.4 |
| 00:38:11 | 20.1 |
| 01:38:11 | 20 |
| 02:38:11 | 20 |
| 03:38:11 | 20.2 |
| 04:38:11 | 20.2 |
| 05:38:11 | 21.3 |
| 06:38:11 | 22.7 |
| 07:38:11 | 25 |
| 08:38:11 | 29.9 |
| 09:38:11 | 28.5 |
| 10:38:11 | 31.6 |
| 11:38:11 | 31.7 |
| 12:38:11 | 32.1 |
| 13:38:11 | 32.2 |
| 14:38:11 | 31.5 |
| 15:38:11 | 28 |
| 16:38:11 | 31.4 |
| 17:38:11 | 27.3 |
| 18:38:11 | 24.7 |
| 19:38:11 | 23.2 |
| 20:38:11 | 22.6 |
| 21:38:11 | 22.1 |
