## Supplementary material for "Elucidating cryptic sympatric speciation in terrestrial orchids": S3

**Appendix 3.** Summary of results of maximum parsimony analyses of *Liparis* (Orchidaceae, Epidendroideae) and outgroups.

| Parameters | ITS | *matK* | Combined |
| --- | --- | --- | --- |
| Characters in matrix | 682 | 1532 | 2219 |
| Variable characters | 446 | 421 | 867 |
| Phylogenetically informative characters | 376 | 238 | 614 |
| Steps | 2046 | 679 | 2755 |
| Most parsimonious trees | 28 | 669 | 18 |
| Consistency Index | 0.41 | 0.72 | 0.48 |
| Retention Index | 0.76 | 0.88 | 0.78 |
