## Supplementary material for "Elucidating cryptic sympatric speciation in terrestrial orchids": S4

P A U P *

Version	4.0b10	for	32-bit	Microsoft	Windows Sat	Mar	18	15:47:01	2017

-----------------------------NOTICE-----------------------------

This	is	a	beta-test	version.	Please	report	any	crashes, apparent	calculation	errors, or	other	anomalous	results. There are no restrictions on publication of results obtained with	this	version, but	you	should	check	the	WWW	site frequently	for	bug	announcements	and/or	updated	versions.

See	the	README	file	on	the	distribution	media	for	details.

----------------------------------------------------------------

Processing	of	file	"E:\Artigos\Liparis	\Liparis	ITS\Liparis	ITS alinhamento_NEX.nex"	begins...

Data	read	in	DNA	format

Data	matrix	has	69	taxa, 682	characters Valid	character-state	symbols: ACGT Missing	data	identified	by	'?'

Gaps identified by '-' "Equate"	macros	in	effect:

R,r	==>	{AG} Y,y	==>	{CT} M,m	==>	{AC} K,k	==>	{GT} S,s	==>	{CG} W,w	==>	{AT} H,h	==>	{ACT} B,b	==>	{CGT} V,v	==>	{ACG} D,d	==>	{AGT} N,n	==>	{ACGT}

Warning! The	following	names	have	been	assigned	to	more	than	one	taxon:

>Liparisnervosa, >Malaxisacuminata

Processing	of	file	"E:\Artigos\Liparis	\Liparis	ITS\Liparis	ITS alinhamento_NEX.nex"	completed.

Heuristic	search	settings: Optimality	criterion	=	parsimony

Character-status	summary: Of	682	total	characters:

All	characters	are	of	type	'unord' All	characters	have	equal	weight

236 characters are constant

70	variable	characters	are	parsimony-uninformative Number	of	parsimony-informative	characters	=	376

Gaps	are	treated	as	"missing"

Multistate	taxa	interpreted	as	uncertainty Starting	tree(s)	obtained	via	stepwise	addition

Addition	sequence: simple	(reference	taxon	=	>Eriaajavanica) Number	of	trees	held	at	each	step	during	stepwise	addition	=	1 Branch-swapping	algorithm: tree-bisection-reconnection	(TBR) Steepest	descent	option	not	in	effect

Initial	'MaxTrees'	setting	=	100

Branches	collapsed	(creating	polytomies)	if	maximum	branch	length	is	zero 'MulTrees'	option	in	effect

Topological	constraints	not	enforced Trees are unrooted

Heuristic	search	completed

Total	number	of	rearrangements	tried	=	3536444 Score of best tree(s) found = 1925

Number	of	trees	retained	=	28 Time	used	=	3.73	sec

Strict	consensus	of	28	trees:

/------------------------------------------------------------	>Eriaajavanica

|| /----- >Acanthophippium

| /---+

| | \----- >Acanthephippium

+--------------------------------------------------+

| \--------- >Collabiumsimple

|

| /--------- >Dieniacylindros

| |

| /----+ /----- >Lipariskoreana

| | \---+

| | \----- >Liparisliliifol

| |

| /---+ /----- >Lipariskumokiri

| | | /---+

| | | | \----- >Liparisfujisane

| /----+ \----+

| | | \--------- >Liparisjaponica

| | |

| /----+ \------------------ >Liparisloeselii

| | |

| /---+ \----------------------- >Liparispauliana

| | |

| /----+ \---------------------------- >Liparisclypeolu

| | |

| /-------------+ \-------------------------------- >Liparisauricula

| | |

| | \------------------------------------- >Lipariskrameri

| |

| | /----- >Crepidiumresupi

| | /---+

| | | \----- >Crepidiumacumin

| | |

| | /--------+--------- >Malaxisacuminat

| | | |

| | | \--------- >Malaxisacuminat

| | |

| | | /----- >Malaxispunctata

| | | /---+

| | | | \----- >Malaxisoculata

| | /----+ /----+

| | | | | \--------- >Malaxishahajime

| | | | |

| | | | +-------------- >Malaxistaurina

| | | | |

| | | | | /----- >Malaxisresupina

| | | \---+--------+

| /---+ /----+ | \----- >Malaxisperakens

| | | | | |

| | | | | +-------------- >Malaxismetallic

| | | | | |

| | | | | \-------------- >Malaxisbreviden

| | | | |

| | | /---+ \----------------------- >Malaxisophrydis

| | | | |

| | | | | /----- >Liparisnervosa

| | | | | /---+

| | | | | | \----- >Liparisnervosa

| | | /----+ | |

| | | | | \------------------+--------- >Liparis sp. “cryptic”

| | | | | |

| | | | | \--------- >Liparis sp. “cryptic”

| | | /----+ |

| | | | | \-------------------------------- >Liparisnervosa

| | | | |

| | | /---+ \------------------------------------- >Liparisformosan

| | | | |

| | | | | /----- >Liparispingxian

| | \----+ \------------------------------------+

| | | \----- >Liparislayardii

| | |

| | \---------------------------------------------- >Liparisrheedei

| |

| | /----- >Lipariscondylob

| | /---+

| | | \----- >Liparisviridifl

| | /----+

| | | \--------- >Liparislatifoli

| | /-------------+

| | | \-------------- >Liparistruncico

\----+	|

| | /----- >Liparisbalansae

| | /---+

| | | \----- >Liparisstrickla

| | /----+

| /---+ | \--------- >Lipariscf.dista

| | | /---+

| | | | \-------------- >Liparisterrestr

| | | /----+

| | | | \------------------ >Lipariscaespito

| | | |

| | \----+ /----- >Liparisnugentae

| | | /---+

| /-------------+ | | \----- >Liparisbracteat

| | | \-------------+

| | | \--------- >Liparissula

| | |

| | | /----- >Liparisanophele

| | | /--------+

| | | | \----- >Liparispandurat

| | | |

| | \-----------------+ /----- >Liparisbrunnesc

| | | /---+

| | | | \----- >Liparisgibbosa

| | \----+

| | \--------- >Liparisdisticha

| |

| | /----- >Oberonianeocale

| | /---+

| | | \----- >Oberoniajaponic

| | /----+

| | | \--------- >Oberoniasetifer

| | /---+

\--------+	|	\--------------	>Oberoniapachyra

| |

| /----+ /----- >Oberoniarecurva

| | | /---+

| | | | \----- >Oberoniaensifor

| | \--------+

| /----+ \--------- >Oberoniabrunoni

| | |

| | | /----- >Oberoniairidifo

| /---+ \-----------------+

| | | \----- >Oberoniamucrona

| | |

| /----+ \---------------------------- >Oberoniafalcone

| | |

| | \-------------------------------- >Oberoniakwangsi

| /----+

| | | /--------- >Oberoniawappean

| | | |

| | \---------------------------+ /----- >Oberoniaequitan

\---+	\---+

| \----- >Oberoniapadange

|

\------------------------------------------	>Oberoniahelioph

Bootstrap	method	with	heuristic	search: Number	of	bootstrap	replicates	=	100 Starting	seed	=	2066359563

Optimality	criterion	=	parsimony Character-status	summary:

Of	682	total	characters:

All	characters	are	of	type	'unord' All	characters	have	equal	weight

236 characters are constant

70	variable	characters	are	parsimony-uninformative Number	of	parsimony-informative	characters	=	376

Gaps	are	treated	as	"missing"

Multistate	taxa	interpreted	as	uncertainty Starting	tree(s)	obtained	via	stepwise	addition

Addition	sequence: simple	(reference	taxon	=	>Eriaajavanica) Number	of	trees	held	at	each	step	during	stepwise	addition	=	1 Branch-swapping	algorithm: tree-bisection-reconnection	(TBR) Steepest	descent	option	not	in	effect

Initial	'MaxTrees'	setting	=	100

Branches	collapsed	(creating	polytomies)	if	maximum	branch	length	is	zero 'MulTrees'	option	in	effect

Topological	constraints	not	enforced Trees are unrooted

100	bootstrap	replicates	completed

Note: Effectiveness	of	search	may	have	been	diminished	due	to	tree-buffer overflow.

Time	used	=	01:41:20.0

Bootstrap	50%	majority-rule	consensus	tree

/--------------------------------------------------------	>Eriaajavanica(1)

|

| /---- >Acanthophippium(2)

| /100-+

| | \---- >Acanthephippium(3)

+----------------------57----------------------+

| \|  \| | \--------- | >Collabiumsimple(4) |
| --- | --- | --- |
| \| | /--------- | >Dieniacylindros(5) |
| \| | \| |  |
| \| | \| /---- | >Lipariskoreana(6) |
| \| | \| \| |  |
| \| | \| +---- | >Liparisliliifol(7) |
| \| | \| \| |  |
| \| | \| +---- | >Lipariskumokiri(36) |
| \| | \| \| |  |
| \| | +-53-+---- | >Liparisjaponica(41) |
| \| | \| \| |  |
| \| | /-----------------99------------------+ +---- | >Liparisloeselii(42) |
| \| | \| \| \| |  |
| \| | \| \| +---- | >Liparispauliana(43) |
| \| | \| \| \| |  |
| \| | \| \| \---- | >Liparisfujisane(52) |
| \| | \| \| |  |
| \| | \| +--------- | >Liparisauricula(27) |
| \| | \| \| |  |
| \| | \| +--------- | >Lipariskrameri(40) |
| \| | \| \| |  |
| \| | \| \--------- | >Liparisclypeolu(53) |
| \| | \| |  |
| \| | \| /---- | >Crepidiumresupi(9) |
| \| | \| /-----72-----+ |  |
| \| | \| \| \---- | >Crepidiumacumin(10) |
| \| | \| \| |  |
| \| | \| \| /---- | >Malaxisacuminat(59) |
| \| | \| +-----71-----+ |  |
| \| | \| \| \---- | >Malaxisacuminat(65) |
| \| | \| \| |  |
| \| | \| \| /---- | >Malaxispunctata(60) |
| \| | \| /-51-+ /-70-+ |  |
| \| | \| \| \| \| \---- | >Malaxisoculata(67) |
| \| | \| \| \| /67-+ |  |
| \| | \| \| \| \| \--------- | >Malaxishahajime(68) |
| \| | \| \| \| \| |  |
| \| | /-86-+ \| \| \| /---- | >Malaxistaurina(61) |
| \| | \| \| \| \| +---52---+ |  |
| \| | \| \| \| \| \| \---- | >Malaxisbreviden(69) |
| \| | \| \| /53-+ \74-+ |  |
| \| | \| \| \| \| \| /---- | >Malaxisresupina(63) |
| \| | \| \| \| \| +---73---+ |  |
| \| | \| \| \| \| \| \---- | >Malaxisperakens(66) |
| \| | \| \| \| \| \| |  |
| \| | \| \| \| \| \------------- | >Malaxismetallic(64) |
| \| | \| \| \| \| |  |
| \| | \| \| /63-+ \---------------------- | >Malaxisophrydis(62) |

| | | | |

| | | | | /---- >Liparisnervosa(48)

| | | | | /-87-+

| | | | | | \---- >Liparisnervosa(51)

| | | /61-+ | |

| | | | | \------100-------+--------- >Liparis sp. “cryptic” (49)

| | | | | |

| | | | | \--------- >Liparis sp. “cryptic” (50)

| | | /-77-+ |

| | | | | \------------------------------ >Liparisnervosa(47)

| | | | |

| | | /88-+ \---------------------------------- >Liparisformosan(54)

| | | | |

| | | | | /---- >Liparispingxian(29)

| | \100+ \----------------99----------------+

| | | \---- >Liparislayardii(30)

| | |

| | \------------------------------------------- >Liparisrheedei(55)

| |

| | /---- >Lipariscondylob(8)

\100+	/100-+

| | \---- >Liparisviridifl(56)

| /100+

| | \--------- >Liparislatifoli(58)

| /-------64-------+

| | \------------- >Liparistruncico(35)

| |

| | /---- >Oberonianeocale(11)

| | /100-+

| | | \---- >Oberoniajaponic(14)

| | /69-+

| | | \--------- >Oberoniasetifer(25)

| | /51-+

| | | \------------- >Oberoniapachyra(17)

| | |

| | | /---- >Oberoniairidifo(12)

| | +----100-----+

| | | \---- >Oberoniamucrona(18)

| | |

| | +----------------- >Oberoniakwangsi(15)

| | /-89-+

| | | | /---- >Oberoniarecurva(16)

| | | | /-61-+

| | | | | \---- >Oberoniaensifor(20)

| | | +--65---+

| | | | \--------- >Oberoniabrunoni(21)

| | /56-+ |

| | | | \----------------- >Oberoniafalcone(19)

| | | |

| | | | /--------- >Oberoniawappean(22)

| | | | |

\---------97----------+100+	\-----50-----+	/----	>Oberoniaequitan(23)

| | \100-+

| | \---- >Oberoniapadange(24)

| |

| \-------------------------- >Oberoniahelioph(13)

|

| /---- >Liparisbalansae(26)

| /-74-+

| | \---- >Liparisstrickla(34)

| |

| +--------- >Lipariscaespito(32)

+---------96---------+

| +--------- >Lipariscf.dista(39)

| |

| \--------- >Liparisterrestr(57)

|

| /---- >Liparisnugentae(28)

| /-93-+

| | \---- >Liparisbracteat(46)

+---------71---------+

| \--------- >Liparissula(44)

|

| /---- >Liparisanophele(31)

| /---97---+

| | \---- >Liparispandurat(45)

| |

\-------97-------+	/----	>Liparisbrunnesc(33)

| /-64-+

| | \---- >Liparisgibbosa(38)

\64-+

\---------	>Liparisdisticha(37)
